# Spatial and temporal control of norovirus protease activity is determined by polyprotein processing and intermolecular interactions within the viral replication complex

**DOI:** 10.1101/175463

**Authors:** Edward Emmott, Alexis de Rougemont, Jürgen Haas, Ian Goodfellow

**Affiliations:** Division of Virology, Department of Pathology, University of Cambridge, Addenbrookes Hospital, Hills Road, Cambridge, UK; National Reference Centre for Gastroenteritis Viruses, Labology of Biology and Pathology, University Hospital Dijon Bourgogne, Dijon, France; AgroSup Dijon, PAM UMR A 02.102 Bourgogne Franche-Comte University, Dijon, France; Division of Infection and Pathway Medicine, University of Edinburgh Medical School, Edinburgh, UK

## Abstract

Norovirus infections are a major cause of acute viral gastroenteritis and a significant burden to human health globally. A vital process for norovirus replication is the processing of the nonstructural polyprotein, by an internal protease, into the necessary viral components required to form the viral replication complex. This cleavage occurs at different rates resulting in the accumulation of stable precursor forms. In this report, we characterized how precursor forms of the norovirus protease accumulate during infection. Using stable forms of the protease precursors we demonstrated that these are all proteolytically active *in vitro*, but that when expressed in cells, activity is determined by both substrate and protease localization. Whilst all precursors could cleave a replication complex-associated substrate, only a subset of precursors lacking NS4 were capable of efficiently cleaving a cytoplasmic substrate. For the first time, the full range of protein-protein interactions between murine and human norovirus proteins were mapped by LUMIER assay, with conserved interactions between replication complex members, modifying the localization of a subset of precursors. Finally, we demonstrate that re-targeting of a poorly cleaved artificial cytoplasmic substrate to the replication complex is sufficient to permit efficient cleavage in the context of norovirus infection. This offers a model for how norovirus can regulate the timing of substrate cleavage throughout the replication cycle. The norovirus protease represents a key target in the search for effective antiviral treatments for norovirus infection. An improved understanding of protease function and regulation, as well as identification of interactions between the other non-structural proteins, offers new avenues for antiviral drug design.

## Introduction

Noroviruses are the causative agent of winter vomiting disease, and following the introduction of the rotavirus vaccine, a leading cause of viral gastroenteritis worldwide. Whilst models for human norovirus infection have recently been developed (1, 2), due to the availability of efficient cell culture and robust reverse genetics systems, murine norovirus (MNV) has been widely used to dissect the molecular details of viral translation and replication (3–5).

Noroviruses are small, non-enveloped positive sense RNA viruses (+ssRNA) forming a genus within the *Caliciviridae* family. As observed with many other+ssRNA viruses, noroviruses generate the majority of the viral replicase components by cleavage of a long polyprotein by the viral protease NS6 (Figure 1A) (6). A key benefit of this strategy is that due to differing rates of cleavage, polyprotein processing generates both the fully processed products and a range of stable precursor intermediates (6, 7). The role of polyprotein precursors in viral replication have been studied extensively for the *Picornaviridae* which share similarities in their polyprotein organisation (8). For noroviruses however, the sole precursor to be examined to date in any detail is the NS6-NS7 precursor (also known as Pro-Pol, or 3CD) (9, 10). Cleavage of the NS6-7 junction is known to be essential for norovirus viability and the NS6-NS7 precursor retains both protease and polymerase activities (9, 10). In contrast, for some members of the *Caliciviridae*, whilst a cleavage site between NS6 and NS7 is present, this precursor remains uncleaved (11).

**Figure 1.**
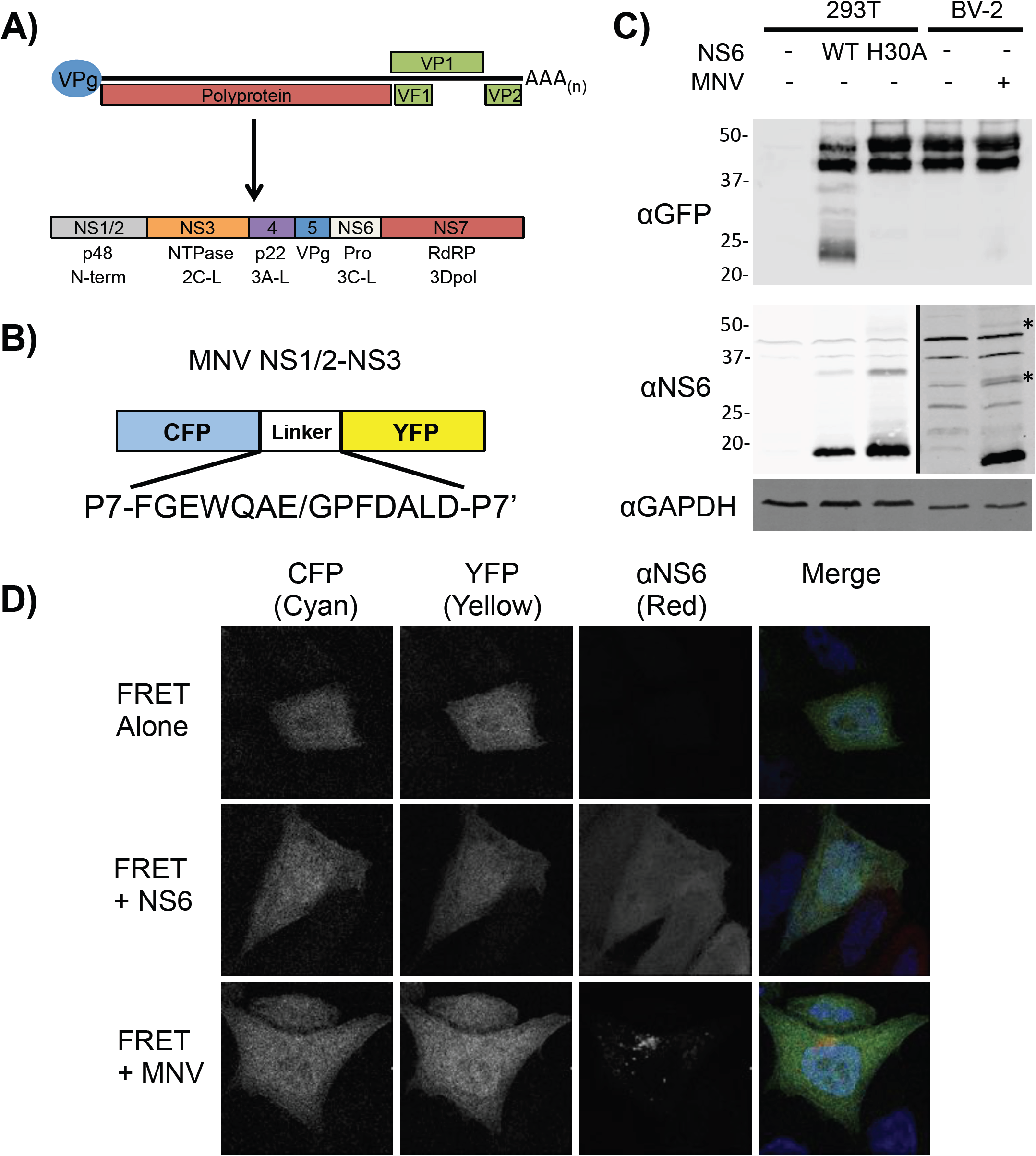
Norovirus protease produced during authentic viral replication inefficiently cleaves a cytoplasmic-localized protease substrate. A) Genome schematic of the murine norovirus genome highlighting the major open reading frames and the components of the polyprotein. B) Schematic of the CFP-YFP FRET sensor highlighting the NS1/2-NS3 boundary cleavage site. C) Western blot analysis of HEK-293T cells co-transfected with the FRET sensor and either wild-type or catalytically inactive protease (WT and H30A respectively), or BV-2 cells electroporated with *in vitro* transcribed RNA encoding the FRET sensor and subsequently infected at MOI 10 TCID_50_/cell with MNV for 9h. Higher molecular weight NS6 containing isoforms produced during viral infection are indicated with asterisks. Note that the left and right-hand regions of the NS6 western blot were imaged at different intensities to compensate for the lower NS6 signal intensity in infected cells. D) Confocal microscopy of HeLa-CD300lf cells transfected with the FRET sensor and either co-transfected with an NS6 expression construct (NS6) and harvested at 18h post-transfection, or infected with MNV (+MNV) at a MOI of 10 TCID_50_/cell at 6h post-transfection, and harvested at 12h post-infection. The FRET sensor was visualized in the CFP and YFP channels, the protease with antisera against NS6, and nuclei stained with DAPI.

Precursor protein functions are often distinct from those of the fully cleaved mature products (12). The rate of precursor cleavage and accumulation can also regulate key stages of the viral replication cycle; for example the cleavage of alphavirus non-structural proteins from the P123 and P1234 polyproteins provides temporal regulation of sense and anti-sense RNA transcript synthesis (13). In addition to cleaving viral substrates, viral proteases also target cellular proteins. We recently identified the host protein poly(A) binding protein (PABP) as a target of the norovirus protease, particularly at late stages in the viral life cycle. The cleavage of PABP forms part of a viral strategy to inhibit host translation, and therefore the production of interferon-stimulated genes (14). However the means by which PABP cleavage was restricted to late times post-infection was unclear.

Processing of the norovirus polyprotein by NS6 results in the production of six proteins; NS1/2, NS3, NS4, NS5, NS6 and NS7 (15) (Figure 1). NS5 functions as both a protein primer for RNA synthesis, as well as permitting translation of the viral genome through interactions with the host translation initiation complex (16, 17). NS6 (also known as Pro, 3C-like) and NS7 (3DPol, RdRP) encode the 3C-like protease and RNA-dependent RNA polymerase respectively. The N-terminal 3 proteins are less well characterized but are known to localize to cytoplasmic membranes. NS1/2 localises to the endoplasmic reticulum (18), and is implicated in viral persistence (19). NS3 has NTPase (20) and predicted RNA helicase activities, and was recently shown to interact with cholesterol-rich membranes and cytoskeletal proteins (21). NS4, an orthologue of the picornavirus 3A protein, is localized to endosomes (18).

In this study we identify a mechanism by which noroviruses regulate viral protease cleavage of substrates during the viral replication cycle. We demonstrate how polyprotein processing controls protease localization and as a result, substrate accessibility, providing a mechanism for temporal regulation of protease cleavage of substrates. Furthermore, we have demonstrated that a comprehensive network of protein-protein interactions occur between the viral replicase proteins which also regulates protease and precursor localization. Finally we demonstrate that targeting an artificial substrate to the viral replication complex allows cleavage by authentic viral protease produced during infection. As the polyprotein includes key drug targets including the protease and polymerase (22, 23), an improved understanding of how these proteins function and are regulated throughout the viral life cycle may aid drug design and targeting.

## Results

### The norovirus protease cleaves a cytoplasmic FRET substrate upon transfection but not in the context of infection

We sought to examine whether norovirus protease precursors play an important role in regulating substrate cleavage. To this aim, we compared the ability of the NS6 protease to cleave an artificial substrate when produced either as a fully processed form or produced during authentic viral replication. We have previously reported the generation of a FRET sensor for assessing norovirus protease activity in live cells (7). This FRET sensor consists of CFP fused to YFP via a short linker region consisting of the P7 to P7’ region of the MNV NS1/2-3 cleavage site (Figure 1B). Whilst the expression of the mature protease resulted in efficient cleavage of the sensor, recapitulating our previous results, protease produced during viral infection showed extremely poor cleavage of the FRET sensor (Figure 1C). Western blotting for NS6 protein confirmed the presence of higher molecular weight isoforms of NS6 during infection (Figure 1C). These higher molecular weight proteins likely represent precursor forms of the protease, that we and others have observed previously (6, 7).

To examine whether poor cleavage of the FRET substrate during infection was due to a mismatch between substrate and protease localization, the localization of the protease and sensor was examined following transfection or infection of HeLa cells expressing the recently identified MNV receptor CD300lf (24, 25). Confocal microscopy revealed that the FRET sensor displayed diffuse nuclear and cytoplasmic localization, in both transfected and infected cells (Figure 1D). Whilst NS6 produced following transfection showed the same diffuse localization as the FRET sensor, in infected cells NS6 was located primarily to a large perinuclear puntca, consistent with previous observations (17) and most likely reflecting the position of the viral replication complex (RC). Given that the NS6 antisera recognizes both mature and precursor forms of NS6, the localization pattern likely represents an averaged localization for both mature NS6 and NS6-containing precursors. These data suggest that substrate cleavage during infection is likely impacted by precursor and substrate localisation, and that the context in which NS6 is expressed may affect the relative efficiency with which a substrate is cleaved.

### Targeting a cytoplasmic substrate to the replication complex permits efficient cleavage by NS6

To determine if poor substrate-protease co-localization was responsible for the low levels of cleavage observed during infection we fused the FRET substrate to the C-terminus of NS7 in the context of full-length polyprotein, with either a WT (ORF1-FRET-WT) or a catalytically inactive protease domain (ORF1-FRET-H30A), in order to drive RC localization. Polyprotein processing was unaffected by fusion to the FRET sensor as the relative level of mature NS5 and NS5-containing precursors was similar to those generated from a MNV full-length infectious clone (MNV-FLC-WT, Figure 2A). The FRET sensor was efficiently cleaved in the context of the ORF1-FRET-WT construct, producing a ∼70kDa NS7-CFP fusion product which was increased in mass when expressed in the context of ORF1 with a catalytically inactive protease (ORF1-FRET-H30A) (Figure 2A). We also observed that the fusion of the FRET sensor to ORF1 resulted in RC localization as the CFP signal localized to the perinuclear region where the RC is known to form (Figure 2B). As expected, the RC localization of the YFP component of the FRET sensor was more pronounced when expressed in the context of ORF1 with a catalytically inactive protease (ORF1-FRET-H30A, Figure 2B). These data confirm that targeting of an artificial protease substrate to the RC allows for efficient cleavage of the substrate, adding to our hypothesis that protease localization and/or the context in which the protease is expressed, affects substrate cleavage efficiency.

**Figure 2.**
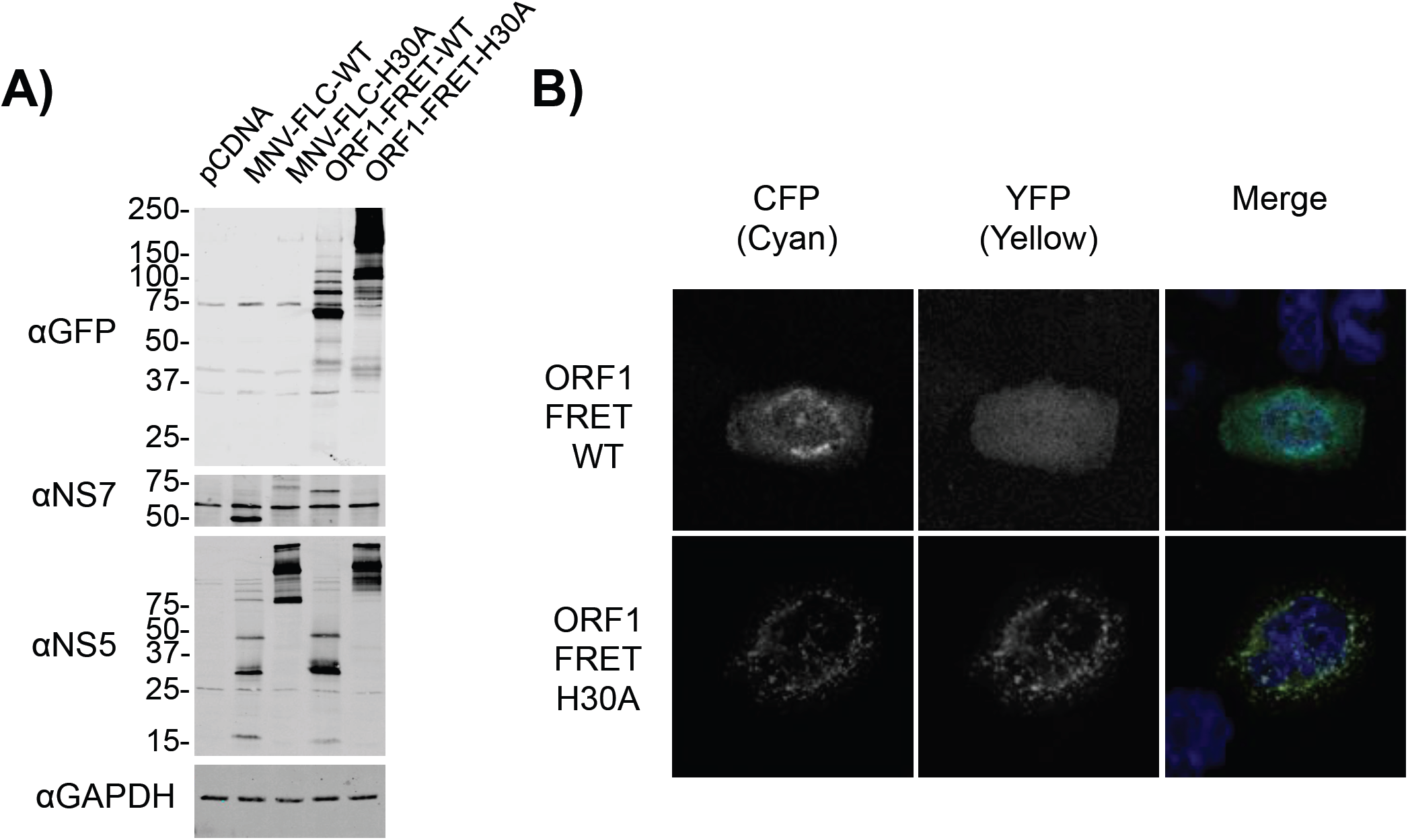
Targeting protease substrates to the norovirus replication complex enhances cleavage by NS6. A) Western blotting of cells BSR-T7 cells transfected with either proteolytically active (MNV-FLC) or inactive (MNV-H30A) full-length infectious MNV clone, or polyprotein fused to the FRET sensor (ORF1-FRET-WT, ORF1-FRET-H30A). Samples were lysed at 18h post-transfection. B) Confocal microscopy of BSR-T7 cells fixed at 18h post-transfection with ORF1-FRET-WT or ORF1-FRET-H30A constructs. The FRET substrate was visualized in the CFP and YFP channels, and nuclei stained with DAPI.

### NS6-containing precursors are abundant and expressed from early times in infection

The processing of the norovirus polyprotein at the five known cleavage sites would, in principle, result in the generation of 21 distinct protein products, of which 10 would be predicted to contain the protease domain (Table S1). To investigate precursor abundance, the presence of precursors during viral replication was examined by western blot (Figure 3). The identity of the observed precursors was assigned based on reactivity with either NS6 or NS5 antisera, predicted molecular weight, comparing migration on SDS-PAGE to a transfected stable form of the individual precursor (Figure S1), and previously published observations (6). NS6-(Figure 3A) and NS5-containing precursors (Figure 3B) were observed as early as 3-4h post-infection, although the predominant NS6 containing precursor seen throughout infection was in fact NS4-6. The majority of the other predicted NS6-containing precursors could be identified including full-length NS1/2-7 (Figure 3A and B).

**Figure 3.**
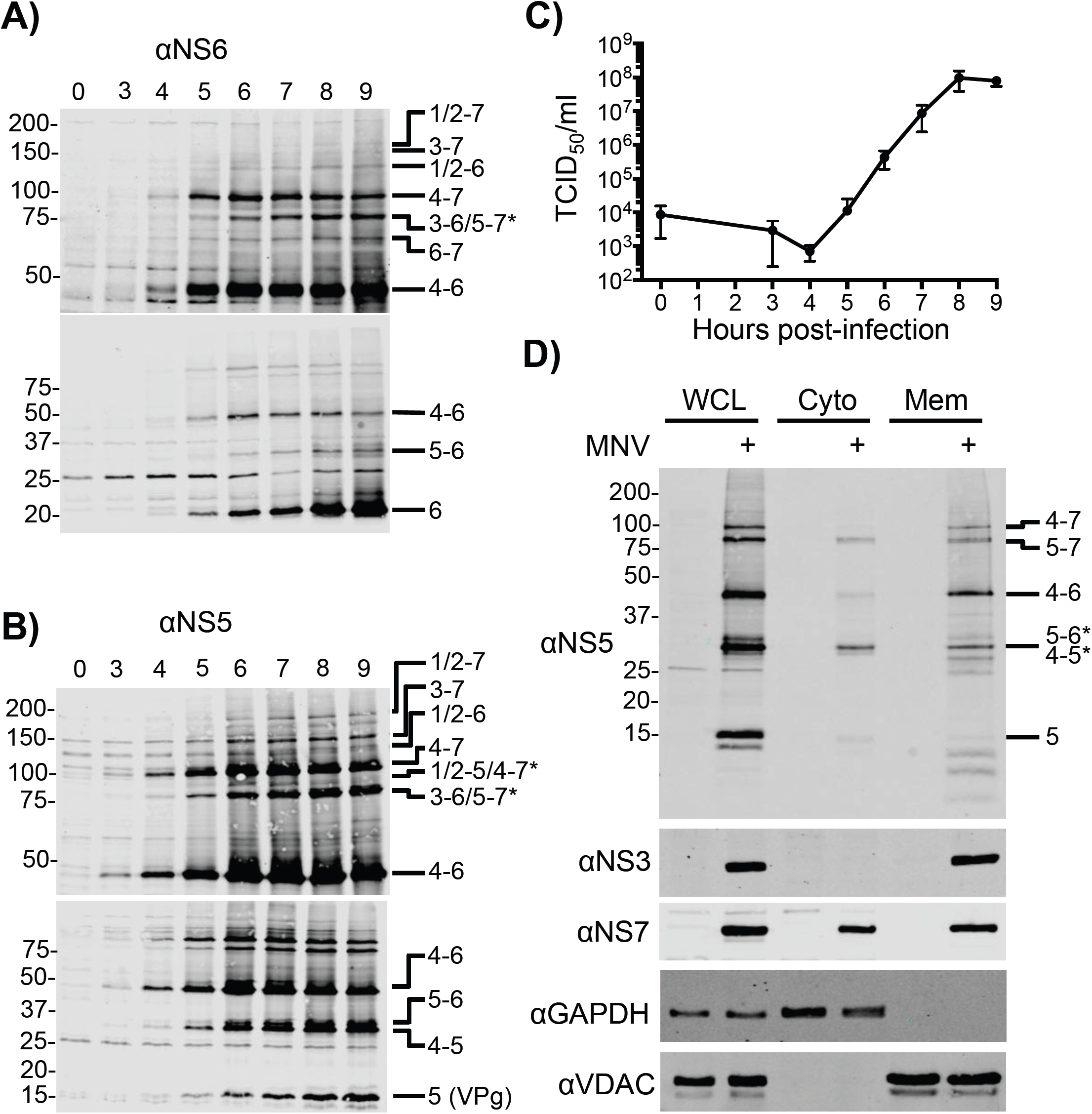
Accumulation of polyprotein precursors during MNV infection. BV-2 cells were infected at a MOI of 10 TCID_50_/cell, and harvested at the indicated times post-infection. Western blot analysis with antisera against A) NS6 or B) NS5/VPg revealed the high levels of precursor proteins present at early stages of infection. To ensure efficient transfer and resolution of the various precursors, samples were separated on both 7.5% (upper) and 17.5% (lower) SDS-PAGE gels prior to transfer and western blot. C) Infectious virus produced across the timecourse was determined by TCID_50_ following freeze-thaw to release intracellular virus. D) To investigate precursor localization, BV-2 cells were infected at MOI 10 and harvested at 9h post-infection where they were subject to fractionation into soluble cytoplasmic, and cytoplasmic membrane-associated fractions. Lysates were blotted for viral and cellular markers as indicated.

Although the NS6-7 precursor was previously suggested to be the major form of NS6 contributing to norovirus polyprotein cleavage based on increased activity *in vitro* compared to mature NS6 (9), our data would indicate that it is produced at relatively low levels. Mature, fully-processed NS6 appears to be the major form of NS6 present over the course of infection. Infectious virus production occurs from 5h post-infection onwards (Figure 3C) at times when the mature form of NS6 and the NS4-6 and NS4-7 precursors are the major forms of NS6 present at detectable levels.

To examine if the localization of NS6 was influenced by the context of the precursors in which it is contained, we used crude fractionation to separate cells into cytoplasmic and membrane-associated fractions (Figure 3D). The distribution of the soluble cytoplasmic and membrane-associated protein marker proteins GAPDH and VDAC confirmed successful fractionation (27). The MNV NS1/2, NS3 and NS4 proteins are known to be membrane associated (18, 21), and our results confirmed NS3 was restricted to the membrane fraction. NS7 associates with the RC, but lacks a predicted trans-membrane domain and could be found in both fractions. The distribution of NS5-containing precursors was used as a surrogate for NS6 due to the significant overlap with NS6-containing precursors and the greater sensitivity of the antisera. High molecular weight precursors containing NS1/2, NS3 or NS4, were absent from the soluble cytoplasmic fraction. In particular the most abundant higher molecular weight precursor (NS4-7) was exclusively isolated in the membrane fraction. Proteins present in the soluble cytoplasmic fraction included fully processed NS5, a protein which may represent NS4/5 or NS5/6, and NS5-7. NS4-6 was present but significantly less abundant in the cytoplasmic than membrane fraction. Some <15kDa proteins were identified in the membrane fraction which may represent modified forms of NS5, or breakdown products. Taken together these data confirm that precursor forms of the norovirus replicase proteins impart distinct patterns of localization on polyprotein constituents, which can differ from their fully processed forms.

### Precursor forms of the norovirus protease are proteolytically active in a cell-free system but show variable activity in cells

As NS6-containing precursors showed variation in their localization in infected cells, we hypothesized that this would also result in variation in their ability to cleave substrates. To test this hypothesis, active, stable forms of these precursors were generated by alanine mutagenesis of the P1 residue (28) at each cleavage site. Of the 21 polyprotein products that can be generated by NS6-mediated polyprotein cleavage, ten contain NS6 (Figure 4A). All protease-containing precursors could be readily expressed *in vitro* and in transfected cells, although to varying degrees (Figure B and C). Expression in both systems yielded proteins of the expected molecular weights (Figure 4A, Table S1). We next examined the ability of each precursor to cleave two different substrates; the ORF1-FRET-H30A substrate, and poly-A-binding protein (PABP), a cellular protein involved in translation that we previously identified as being cleaved by NS6 during norovirus infection (14). When unlabeled *in vitro* translated norovirus protease-containing precursor proteins were incubated with *in vitro* translated ^35^S-methionine labelled PABP, cleavage products were observed (Figure 4D). As expected, cleavage products were not observed upon incubation of PABP with catalytically inactive protease (H30A). It is noteworthy that only a fraction of the PABP produced *in vitro* was cleaved, fitting with our previous observations (14). However, when the protease precursors and FLAG-tagged PABP were co-expressed in cells, a C-terminal PABP cleavage product, detected by virtue of the C-terminal FLAG tag, was only observed when co-expressed with NS6, NS6-, NS5-6 and NS5-7(Figure 4E). The ability of the precursors to cleave a protease substrate that was localized to the viral replication complex was examined by co-transfection of cells with plasmids expressing the precursors and the ORF1-FRET-H30A FRET substrate (Figure 4F). FRET substrate cleavage was observed in all cases except when co-transfected with a catalytically inactive mature protease. In a limited number of cases, in addition to the fully cleaved FRET substrate, additional larger CFP/YFP containing precursors were observed with some protease-containing precursors that were not detected following co-expression with the mature protease. Together these data highlight that the context in which NS6 is expressed and the localization of any given substrate, alters the relative efficiency of substrate cleavage.

**Figure 4.**
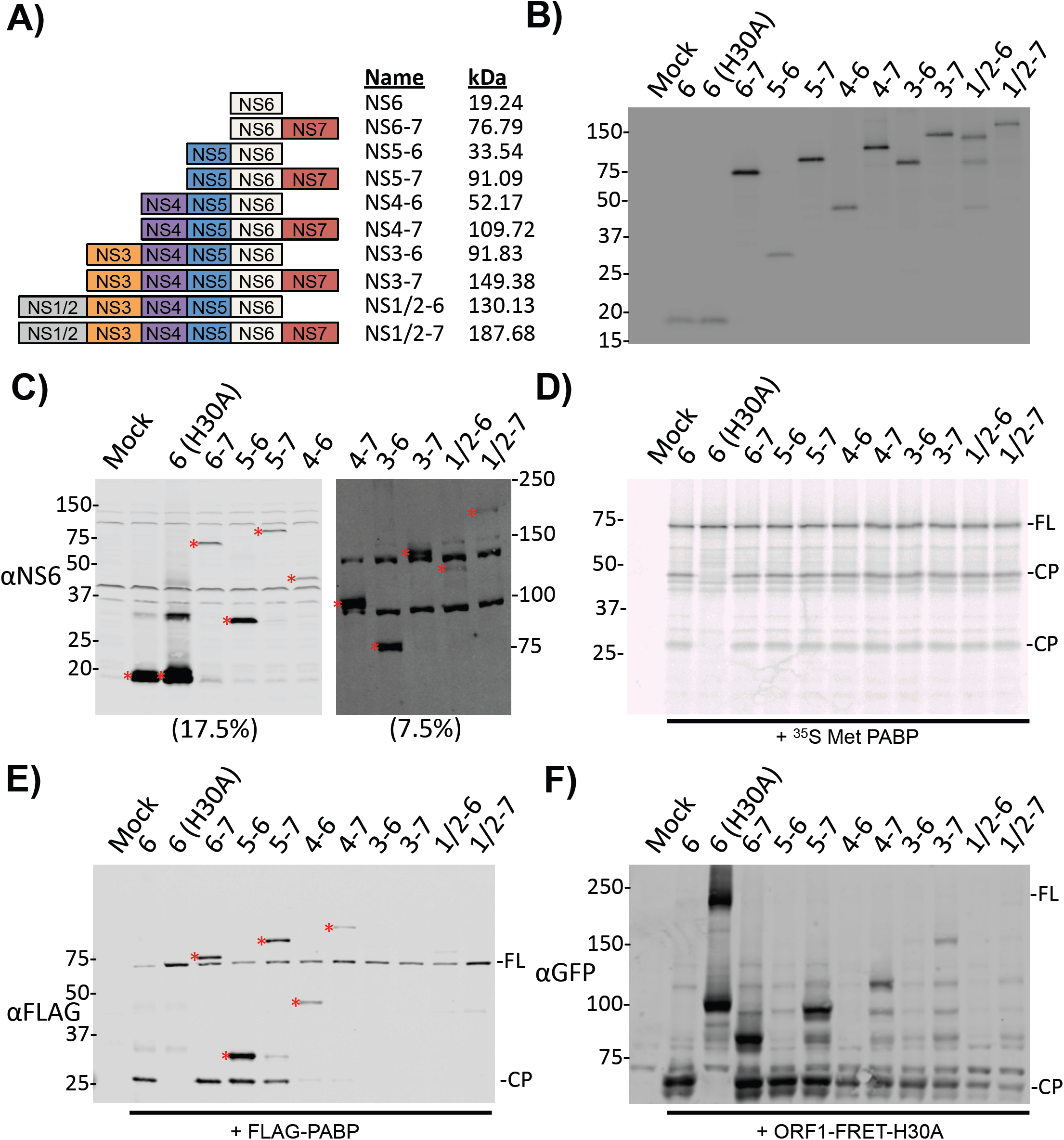
Activity of NS6-containing stable precursors. FLAG-tagged forms of the various protease-containing precursors were generated by mutagenesis of the P1 residue at each cleavage site to an alanine. A H30A mutation within the protease abolished proteolytic activity. A) A schematic illustrating the 10 possible NS6-containing precursors generated by NS6 cleavage of the polyprotein. Protease-containing precursors may all be observed following B) *in vitro* translation of the precursors incorporating ^35^S Met labeling followed by phosphorimaging, or C) Expression in BSR-T7 cells visualized by western blotting. Incubation of unlabeled *in vitro* translated precursors with ^35^S Methionine-labeled substrate demonstrates cleavage of the D) cellular substrate PABP. E) Western blotting of BSR-T7 cells harvested at 18h post-transfection with the precursors and FLAG-PABP reveals only a subset of precursors cleave PABP in cells. FLAG-tagged precursors are also visible on this image and are indicated with an asterisk (*). F) Western blotting of BSR-T7 cells co-expressing ORF1-FRET-H30A as substrate for trans-cleavage by co-expressed protease precursors shows cleavage of this substrate by all precursors at 18h post-transfection.

### NS6 precursor localization is determined by fusion of NS6 with membrane-bound polyprotein components and intra-replication complex interactions

To examine how the context in which NS6 is expressed affects its localization we examined the localization of each precursor in HeLa cells expressing the MNV receptor (CD300lf) by confocal microscopy (Figure 5). The mature fully processed form of NS6 and the NS5-6 precursor showed predominantly cytoplasmic localization, with some nuclear localization. The localization of the other precursors was more varied. NS6-7 also showed some degree of nuclear localization but was largely localized to the cytoplasm. NS5-7 was exclusively localized to the cytoplasm but also formed some cytoplasmic puncta. NS4-6 and NS4-7 showed reticular localization, reminiscent of RC localization. A detailed analysis of the localization of the longer precursors was confounded by the relatively poor expression and toxicity of these precursors following transfection however a limited number of cells could be visualized by confocal microscopy Figure S2. The NS3-containing precursors NS3-6 and NS3-7 localized to small, spherical cytoplasmic membranous structures whereas the NS1/2-6 and NS1/2-7 precursors were largely cytoplasmic, with some cytoplasmic, possibly membrane associated puncta, observed on the case of NS1/2-6. In summary, we found that the precursors containing NS4 and NS3 imparted membrane localization on NS6, consistent with these proteins localizing to the viral RC. These data were also consistent with the poor efficiency with which these precursors cleave PABP (Figure 4E).

**Figure 5.**
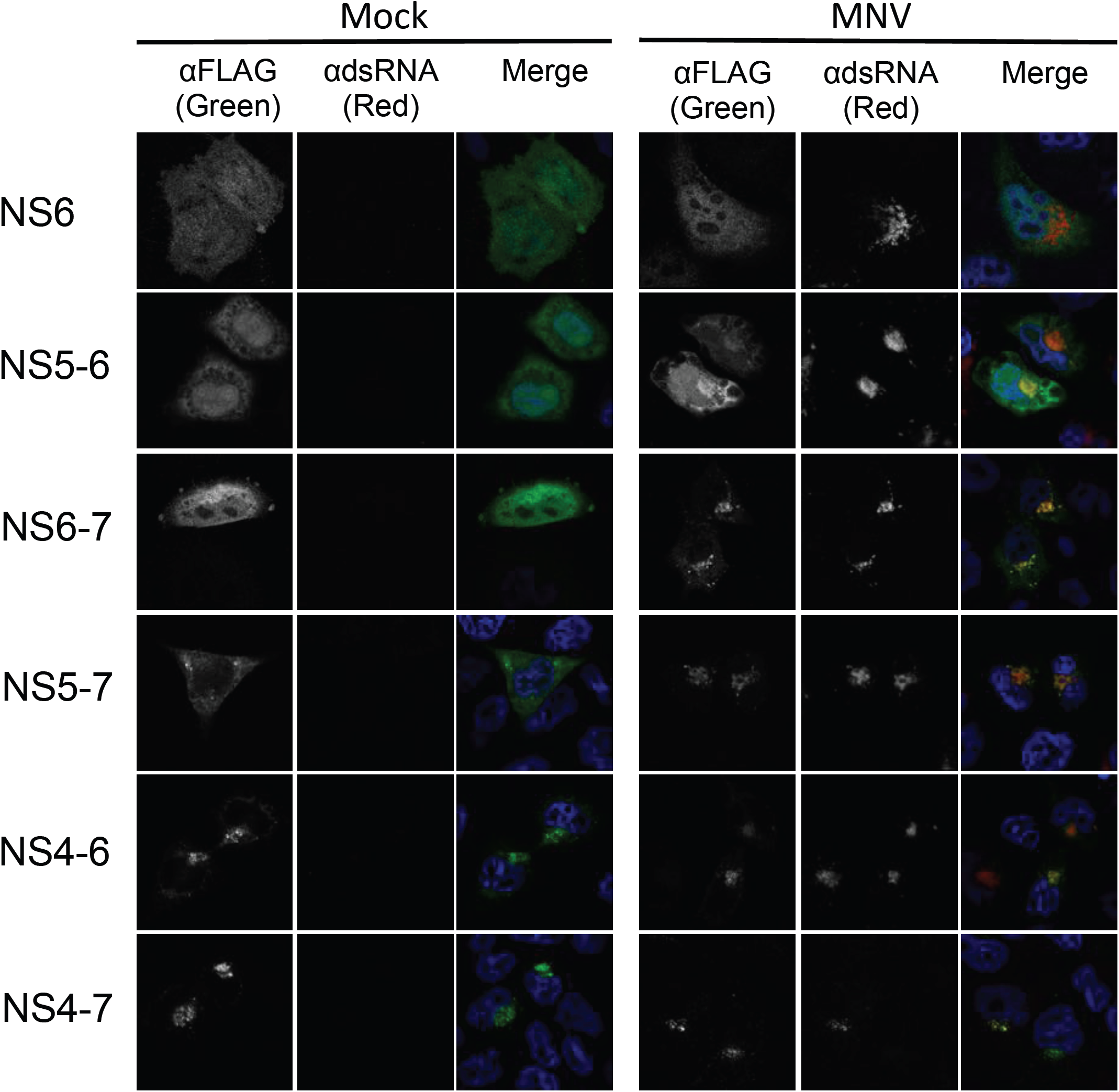
Confocal microscopy of protease precursors in mock or MNV-infected cells. HeLa-CD300lf cells were transfected with the various protease precursor constructs and at 6h post-transfection, mock- or infected with MNV at MOI 10. The cells were fixed and processed for microscopy at 12h post-infection. Nuclei were visualized with DAPI, the precursors with anti-FLAG antisera, and infected cells with anti-dsRNA antisera.

Previous studies indicate that NS7 localizes to the viral RC suggesting in the context of viral infection, suggesting that protein-protein interactions between the viral replicase proteins likely also affects polyprotein localization. To test this hypothesis we examined the impact of viral infection on the localization of each NS6 containing precursor (Figure 5). Whilst the localization of mature NS6, NS4-6 and NS4-7 was not altered upon infection, the NS5-6, NS6-7 and NS5-7 precursors were redistributed to the viral RC. In the case of NS5-6, whilst cytoplasmic and nuclear localization was still visible, there was clear enrichment at the viral RC as evident by colocalization with viral dsRNA. NS6-7 and NS5-7 showed a more pronounced relocalization following infection, with the vast majority of these precursors localized to the RC.

### Identification of protein-protein interactions between norovirus proteins

The relocalization of NS7-containing precursors to the viral RC upon infection highlighted the potential role of interactions between viral components to tether NS6-containing precursors to the viral RC. We therefore examined the network of interactions between norovirus replicase proteins in order to determine which of these might impact on precursor localization. To this aim we used the LUMIER system (29, 30) to map all the interactions between individual MNV proteins (Figure 6 and S3). A similar analysis was also performed for GI.1 and GII.4 human norovirus (Figures S4 and S5) to identify conserved intra-RC protein-protein interactions. Strong self-association was observed with NS1/2, NS4, NS7, and the major and minor capsid proteins VP1 and VP2. The N-terminal components of the polyprotein NS1/2, NS3 and NS4 interacted with each other, all of which have previously been shown to localize to cytoplasmic membranes in order to initiate RC formation for the related feline calicivirus (31). As expected, a robust interaction between the VP1 and VP2 capsid proteins was seen along with VP1-NS7 and NS1/2-NS7 interactions. These latter interactions fit with our previous observations of the impact of VP1 and NS1/2 on polymerase function (32). A interaction was also observed between NS5 and NS7, in keeping with the role of NS5 in serving as a protein primer for RNA synthesis. These data suggest that the redistribution of the NS6 containing precursors NS5-6, NS6-7 and NS5-7 to the viral RC is therefore the likely result of interactions between NS1/2 and NS7, which in the case of NS5-6 could occur via the formation of a NS5-6:NS7:NS1/2 complex.

**Figure 6.**
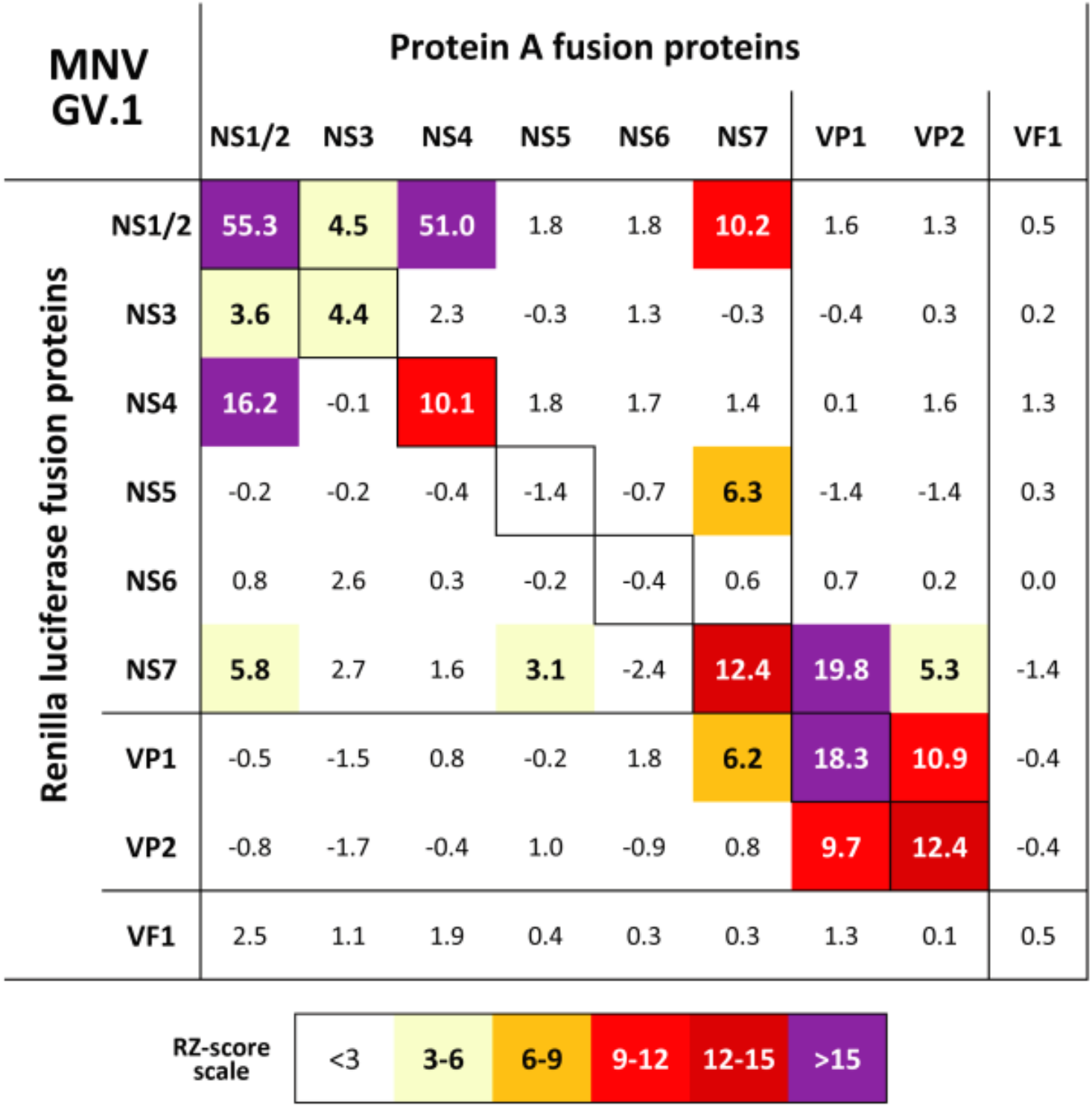
Components of the norovirus replication complex exhibit homo- and heteromeric interactions. HEK-293T cells expressing Protein-A and Renilla luciferase fusions of the various MNV proteins were used for LUMIER analysis to identify protein:protein interactions. The numbers are robust z-scores. Positive protein:protein interactions are coloured by the strength of interaction with weak interactions showing in pale yellow, with the strongest interactions in purple. Protein expression was confirmed by western blotting (Figure S4).

### Fusion to N-terminal replication complex components permits effective cleavage of fused substrates by the norovirus protease *in trans*

To examine the minimum features requires for the correct localization of a substrate to the viral RC and its subsequent cleavage, a range of truncation and deletion mutants in the ORF1-FRET-H30A construct were generated to determine which components were necessary and sufficient for cleavage *in trans* (Figure 7A). Active RC localized viral protease was provided *in trans* by co-expressing the FRET substrates with a MNV full length cDNA clone with either a wild-type (MNV-FLC) or catalytically inactive protease (MNV-H30A).

**Figure 7.**
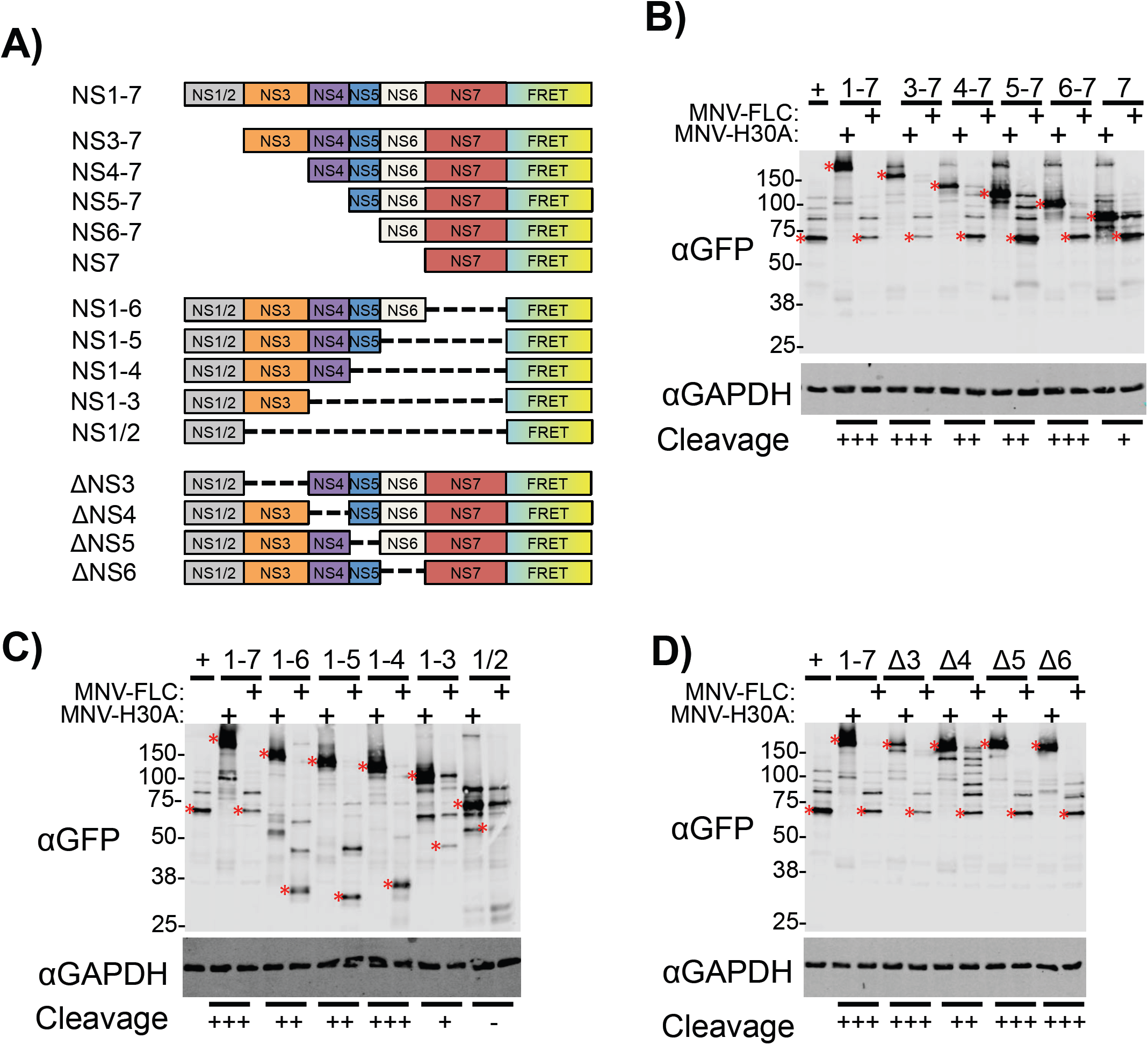
Mutagenesis of polyprotein-FRET fusions reveals the importance of individual polyprotein components for replication complex targeting and substrate cleavage. A) Schematic illustrating the various N-, C- and internal deletion mutants generated from ORF1-FRET-H30A. These mutants were transfected into BSR-T7 cells to function as trans-cleavage substrates. WT or proteolytically-inactive full length clone (MNV-FLC, MNV-H30A) was provided *in trans* to determine cleavage efficiency. Samples were harvested at 18h post-transfection. B) N-terminal, C) C-terminal and D) Internal deletions are shown. Substrate cleavage was assessed using anti-GFP antisera. ORF1-FRET-WT was used as a positive control (+). For clarity the position of either the full length (Mock cells) or fully-processed cleavage product (MNV infected cells) is highlighted with a red asterisk (*).

The N-terminal truncations all displayed some degree of cleavage with the NS7-FRET substrate being cleaved the least efficiently (Figure 7B). Whilst the fusion of NS7 alone to a substrate was sufficient to allow *trans* cleavage, C-terminal truncations demonstrated that NS7 was not essential as the NS1/2-6, NS1/2-5 and NS1/2-4 fusions were all cleaved efficiently (Figure 7C). The NS1/2-3 and NS1/2 fusions showed progressively poorer cleavage of the fused substrate, with negligible cleavage observed with the latter.

Finally, the impact of internal deletions on FRET substrate cleavage was examined (Figure 7D). The ΔNS4 mutant showed a minor defect in cleavage, which may have been due to the chimeric nature of the NS3/5 cleavage site generated by the deletion. Single deletions of NS3, NS4, NS5 and NS6 all permitted efficient cleavage. From this data it is apparent that no single region of the polyprotein was essential for cleavage of a fused substrate, but that several regions of the polyprotein (NS1/2-4 or NS7) appeared sufficient. These regions matched those identified earlier as sufficient for RC localization (Figure 6), either directly, or through interactions with other viral RC components.

### Replication complex localization of FRET substrates permits cleavage by the norovirus protease in the context of infection

Having identified the minimal regions required for RC-targeting of fused substrate by polyprotein and precursors provided *in trans*, we predicted that these fusions would permit the cleavage of said substrate by MNV in the context of infection. The full length ORF1-FRET-H30A fusion and three of the deletion constructs (NS7, NS1/2-4, NS1/2) were chosen for further characterisation.

As anticipated, the ORF1-FRET-H30A fusion was efficiently cleaved during viral infection at both 9h (Figure 8A) and 12h (Figure 8B) post-infection. NS1/2-4-FRET was mostly cleaved at 9h post infection and showed full cleavage by 12h post infection. This NS1/2-4 fusion represented the most efficiently cleaved of the truncated substrates. The NS1/2-FRET fusion showed poor cleavage at 9h post-infection (Figure 8A), consistent with our previous observations (Figure 7C), but cleavage was significantly increased by 12h post infection. The NS7-FRET fusion remained largely uncleaved at both 9h or 12h post-infection (Figure 8A/B). Combined, these data highlight that substrate accessibility and protease localization, may be at least one mechanism by which norovirus temporally regulates the cleavage of any given substrate.

**Figure 8.**
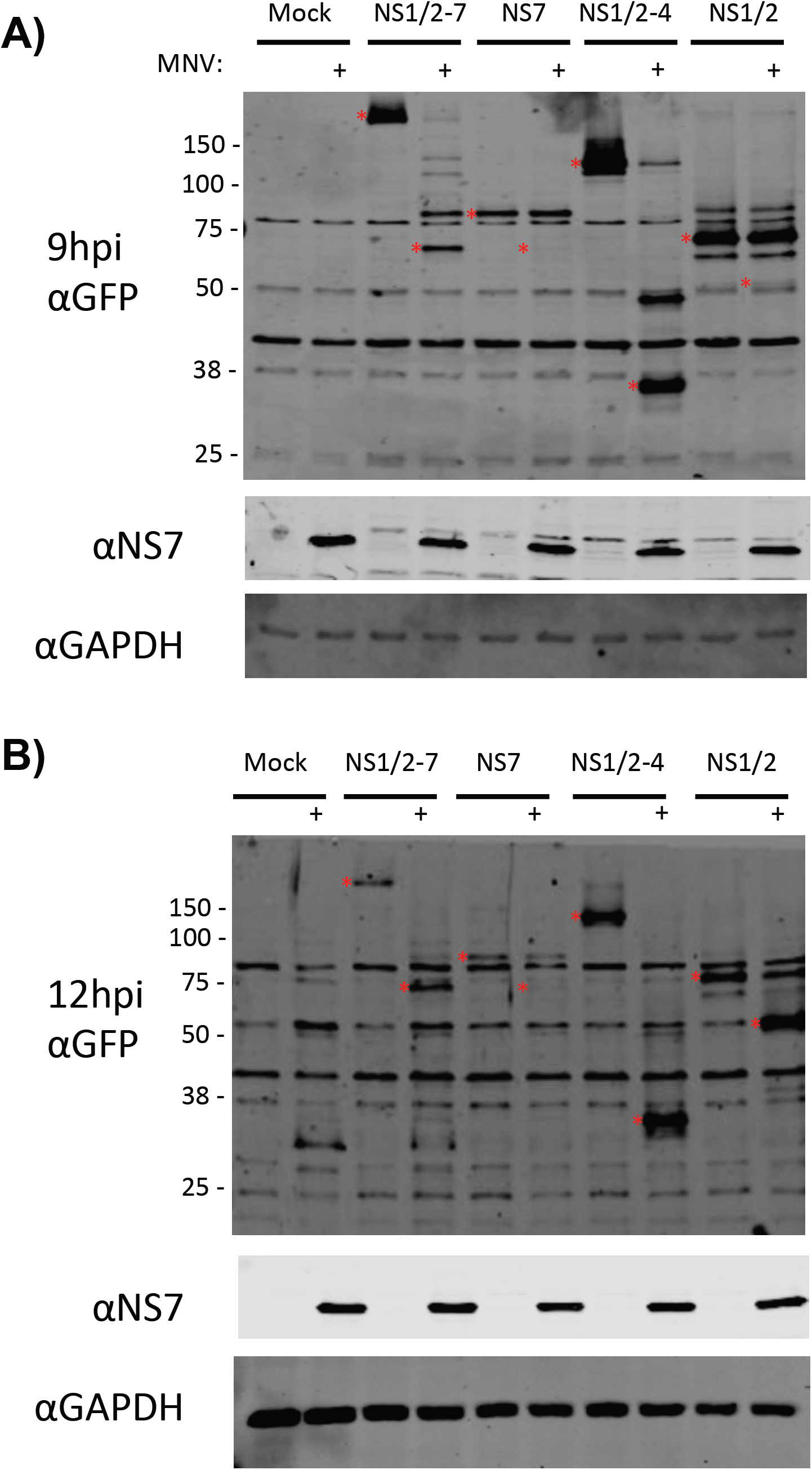
Substrate targeting to the replication complex permits NS6 cleavage in infection. *In vitro* transcribed RNA encoding select truncation mutants of ORF1-FRET-H30A was electroporated into BV-2 cells. At 6h post-electroporation these cells were infected at MOI 10 and lysates prepared at A) 9h or B) 12h post-infection. Trans-cleavage of the substrate was determined by anti-GFP staining, and the presence of active protease provided *in trans* by anti-NS7 staining. For clarity the position of either the full length (Mock cells) or fully-processed cleavage product (MNV infected cells) is highlighted with a red asterisk (*).

## Discussion

The generation of viral replicase components from a single large polyprotein is a mechanism conserved in many +ssRNA viruses including noroviruses. The regulated production of viral precursors, often with distinct functions, enables the expansion of functional coding capacity without a concomitant increase in viral genome size.

In the case of noroviruses, 21 possible precursors can be generated by NS6 cleavage of the polyprotein, ten of which contain the protease. In this study we confirm that all ten of these protease-containing precursors represent active forms of the protease, though spatial sequestration of a number of these precursor forms controls their substrate accessibility. Due to similarity with other caliciviruses, where the protease and polymerase (NS6-7) remain uncleaved, NS6-7 had previously been suggested to represent the dominant form of NS6 proteolytic activity (33). In support of this hypothesis, NS6-7 retains both proteolytic and polymerase activity whilst fused (9, 10), can more efficiently nucleotidylate NS5 than NS7 alone (34), and NS6-7 has higher proteolytic activity than NS6 alone *in vitro* (33). However, our data questions the extent to which this precursor may contribute to these activities as we identified the NS4-6 precursor as the most abundant precursor form produced over the course of an infection. We also found that NS6-7 does not localize directly to the viral RC, but instead requires other viral factors such as NS1/2 for this localization. In addition, we have previously shown that the NS6-7 cleavage site is efficiently cleaved, consistent with the extremely low levels of NS6-7 observed throughout our timecourse experiments (7). Whilst abundant, the diffuse localization of fully cleaved NS6 would result in lower concentrations of this form of the protease both within the replication complex and throughout the cytoplasm as a whole. This would prevent efficient cleavage by NS6 until very late times post-infection. We would therefore hypothesise that NS4-6 represents the form of the protease responsible for the majority of *trans* polyprotein cleavage observed during norovirus infection. The cytoplasmic localization of mature NS6 could provide a mechanism of controlling the levels of viral protein cleavage by expelling excess protease from the RC, and thus avoiding runaway levels of proteolysis as polyprotein abundance increases. In addition, this may allow for the temporal regulation of the cleavage of certain cellular substrates. It should however be noted that whilst this study primarily focuses on *trans*-cleavage by the protease, cis-cleavage most likely plays an important role in norovirus polyprotein processing and will require further research.

In a recent study we identified PABP as a cellular substrate of the norovirus protease (14). Cleavage of PABP at late times post-infection partially accounted for inhibition of host translation and diminished production of innate immune regulatory proteins during MNV infection. Transfection of a non-cleavable form of PABP resulted in reduced in MNV replication (14). The mechanism behind the temporal regulation of PABP cleavage is unknown as mature protease is produced early in infection. From the data presented here, the shorter, non-membrane associated forms of NS6 more effectively cleaved PABP. PABP is a cytoplasmic protein much like the unfused FRET substrate. Whilst the fully-cleaved NS6 is the most abundant form of the protease for much of infection, its diffuse localization would be expected to result in lower concentrations of total protease in the cytoplasm, compared with the replication complex which would also possess the membrane-associated protease precursors. This hypothesis is consistent with the strong replication complex localization observed using anti-NS6 antisera in MNV-infected cells. The concentration of NS6 activity within the RC until late times post-infection may represent the means by which PABP cleavage is temporally regulated, with PABP cleavage only observed once cytoplasmic NS6 concentrations have reached a threshold level.

A network of interactions between viral replication complex proteins, their precursor forms, and a range of cellular proteins is required to establish the formation of a functional viral RC (35, 36). The interactions between the feline calicivirus replicase proteins have been previously investigated by our group using the yeast two-hybrid system (37). In the current study we extended this to include noroviruses, identifying a network of evolutionarily conserved interactions. Surprisingly, we observed no direct interactions with the mature NS6 and any other viral proteins. Further studies are required to determine if this is due to confounding effects of the additional sequences added for the LUMIER assay. However, NS6 has the potential to interact with NS1/2, NS5 and VP1 when part of the NS6-7 precursor, and NS4-6 precursors would also be expected to interact with NS1/2 and NS3 through interactions with the NS4 component of the precursor. Several distinctions between norovirus protein-protein interactions and the previous calicivirus protein interaction study were observed. FCV p76 (equivalent to NS6-7) weakly interacted with the minor capsid protein (37), whereas no interactions between VP2 and the non-structural proteins were observed for the noroviruses. For noroviruses NS1/2 appears to play the key role in binding a number of the non-structural proteins together by interacting with NS3, NS4 and NS7, with FCV p32 playing a similar role (37). However the self interactions of NS3 and NS4 observed in the LUMIER data were not conserved in their FCV p39 and P40 equivalents (37).

Cleavage of a subset of FRET fusion substrates showed some variation between infection and transfection-based *trans*-cleavage studies. NS1/2 fusions, unlike NS1/2-4 fusions, were poorly cleaved following co-transfection with active protease, but were subsequently cleaved, albeit at very late times, in the context of MNV infection. However, the superior cleavage of the NS1/2-4 fusion can be understood in the context of the LUMIER data which highlights interactions between NS1/2, NS3 and NS4. Both substrate and protease expression in the trans-cleavage assay system is driven to high levels by T7-driven expression. In particular, this would be expected to produce significantly more NS1/2-FRET in this system, without a concomitant increase in the NS3 and NS4, an issue not present with the NS1/2-4-FRET substrate. This aberrant stoichiometry between NS1/2, NS3 and NS4 may result in not all of the NS1/2-FRET substrate integrating successfully into the RC. At the lower levels of substrate expression achieved following electroporation into BV-2 cells, a higher proportion of the NS1/2-FRET substrate may be incorporated into the RC, albeit at late times post infection resulting in the cleavage of this substrate at 12h but not 9h post-infection.

The alphaviruses represent the best-characterised example of polyprotein cleavage driving the staging of events in the viral life cycle throughout replication (13). Compared with noroviruses, the alphaviruses have a simpler polyprotein with two alternate forms P123 and P1234 produced by stopcodon readthrough. The successive processing of this polyprotein at the P34, P12 and P23 junctions respectively controls a shift in viral transcription from negative to positive sense transcription (13). Accumulation of the protease, and control of cis- and trans-cleavage activity throughout infection drives these shifts in precursor populations and thus yields a mechanism for temporal regulation of viral replication.

Whilst our current understanding of the roles of norovirus precursor proteins and polyprotein cleavage is less developed than for the alphaviruses, the localization, interactions, and cleavage patterns of the various protease precursors gives rise to a basic mechanistic model for how polyprotein cleavage could regulate some aspects of the norovirus life cycle. Following release of the viral genome has into the cytoplasm, pioneer rounds of translation of the polyprotein from the incoming genome occur. The synthesis of viral negative sense RNA is thought to occur via a *de novo* initiation mechanism that is enhanced by an interaction between the viral capsid protein VP1 and the polymerase NS7 (38–40). The production of viral VPg-linked positive sense RNA can then occur. Analyses of NS5/VPg-linked RNA produced in norovirus infected cells have to date only found fully processed NS5 linked to RNA (16). However, experiments with the distantly-related picornaviruses showed that in cases where the NS5-6 equivalent (3BC) cleavage site was mutated, viable virus was obtained and 3BC could be identified bound to the RNA (41). This suggests that the priming functions of NS5 may not require the fully processed protein. However our group and others have confirmed that the fully processed NS5 is most efficient at binding to eukaryotic initiation factor proteins and therefore translation of viral proteins (16, 42). For example, Leen and colleagues demonstrated that the disruption of NS5-6 cleavage abolished the interaction of NS5 with the eIF4G HEAT domain (42). Regulation of the levels of NS5-containing precursors may therefore provide a mechanism of temporal regulation of viral RNA synthesis, due to the progression in precursor abundance from dominant NS4-6 and NS4-7 forms at early times, to later times when the fully cleaved and shorter precursors (NS4-5, NS5-6) are more abundant. Negative sense RNA synthesis would be anticipated to dominate until sufficient NS5 is present in a suitable (presumably shorter) precursor form to permit sense transcription. In addition this temporal regulation of the NS5 precursor state may also control levels of translation, though how distinct the NS5 cleavage state requirements for efficient translation are from those for positive sense RNA synthesis remains to be fully elucidated.

In conclusion, our study identified a mechanism by which noroviruses may regulate protease activity. This is achieved by both the temporal control of precursor production and the spatial regulation this imparts either directly or indirectly via intra-RC protein-protein interactions with other viral replicase components. Membrane tethering keeps precursors associated with the RC, serving to concentrate their activity and ensure efficient cleavage of viral or RC-associated substrates. At late times shorter forms of the protease lacking either a transmembrane fusion partner such as NS4, or the ability to interact indirectly with the replication complex via NS7 dominate. These forms are no longer sequestered at the RC and are capable of cleaving cytoplasmic substrates. This understanding of which forms of the protease are most active throughout infection, and their regulation may serve to guide future drug design efforts against the protease and other RC components.

## Methods

### Cells and Viruses

Murine BV-2 cells were used for infection experiments (25); human HeLa cells expressing the murine norovirus receptor (HeLa-CD300lf) and hamster BSR-T7 cells were used for transfection and microscopy experiments. Experiments using fowlpox T7-mediated expression of the polyprotein, or polyprotein fusions from the MNV reverse genetics system were performed in BSR-T7 cells as described (43). LUMIER assays were performed in unmodified HEK-293T cells.

All infections were performed with murine norovirus strain CW1 (43). Preparation and titration of virus stocks by TCID_50_ was performed as described (44). All infections were performed at high multiplicity of infection (MOI: 10 TCID_50_/cell). Cells were cultured in DMEM media supplemented with penicillin/streptomycin, 10% serum, and were maintained at 37°C, 5% CO_2_. With the exception of BV-2 cells, transfections were performed in antibiotic-free media using Lipofectamine 2000 (Life Technologies) as per the manufacturer’s recommendations. BV-2 cells were electroporated with *in vitro* transcribed RNAs using the Neon electroporation system (Life Technologies) as described previously (45).

### Plasmids and molecular cloning

The FRET plasmid used was essentially as described in (7). To avoid difficulties in PCR amplification of the FRET sensor (CFP-cleavable linker-YFP) due to high sequence identity between the CFP and YFP coding sequences, this was re-cloned using a modified YFP nucleotide sequence whilst maintaining the original linker and protein sequences (sequence available upon request). The generation of the FLAG-PABP plasmid has been described previously (14).

N-terminally FLAG-tagged stable forms of all the potential NS6 containing precursors were generated in pTriex1.1 (Novagen) using pT7 MNV 3’RZ (43) as a template. In each precursor, all the known cleavage sites had their P1 residue mutated to alanine to inhibit cleavage. Cloning was accomplished by Gibson assembly into the NcoI and XhoI restriction sites of pTriex1.1.

The pT7 MNV 3’RZ plasmid used has been described previously (43), and represents a full-length clone of the MNV CW1 strain (DQ285629), and is referred to as MNV-FLC in the manuscript. The proteolytically inactive form of this plasmid was generated by alanine mutagenesis of His30 within the NS6 sequence (MNV-H30A). The full-length FRET fusion plasmid (ORF1-FRET-WT) and inactive mutant (ORF1-FRET-H30A) were derived from pT7 MNV 3’RZ and had the subgenomic region deleted and the FRET sensor fused to NS7 in place of the NS7 stop codon. A poly(A) tail was included between the stop codon of the FRET sensor and the HDV ribozyme. This permitted expression of this protein by transfection of the plasmid into BSR-T7 cells infected with fowlpox-T7, or by electroporation of *in vitro* transcribed RNA into BV-2 cells.

The various deletion constructs based on ORF1-FRET-H30A were generated by a Gibson assembly approach. N-terminal deletions all incorporated the methionine from NS1/2. C-terminal deletions fused the FRET sensor to the C-terminal-most protein which had its C-terminal P1 residue directly joining the FRET sensor mutated to alanine to prevent off-target cleavage between the C-terminal protein and CFP. For internal deletions, the sequence deleted extended from the P1’ of the N-terminal cleavage site to the P1 of the C-terminal cleavage site of the protein of interest resulting in the production of a chimeric cleavage site. For example, in the NS3 deletion mutant, the cleavage site between NS1/2 and NS4 consists of the P side residues of the NS1/2-NS3 cleavage site and the P’ side residues of the NS3-NS4 cleavage site.

All MNV and genogroup I and II HuNoV genes encoding individual proteins were cloned from MNV-1 (strain CW1) and Norwalk (GI.1) into the Gateway^®^ pDONR207 entry plasmid, and MD145 (GII.4) strains into the Gateway^®^ pDONR221 entry plasmid. N-terminal Renilla reniformis Luciferase (RL) and *S. aureus* protein A (PA) fusion protein plasmids were generated in pcDNA3-RL-GW and pTrex-Dest30-PA expression vectors respectively using the Gateway^®^ Clonase^®^ II enzyme recombination system (Invitrogen). The negative control plasmid consisted of a double protein A fusion protein (PA-PA). Validation of LUMIER assays was performed using positive control plasmids consisting of RL-c-Jun and PA-c-Fos fusion proteins.

### Antibodies

Recombinant NS6 from GI.1, GII.4 and MNV was prepared in E. coli as described previously (46), combined, and used as antigen for preparation of rabbit polyclonal antisera. The resulting sera was affinity purified against MNV NS6. NS3, NS5 (VPg) and NS7 antibodies were prepared in our laboratory previously, and anti-NS5 was affinity purified against Cherry-tagged MNV NS5. Antisera to GAPDH (Ambion), VDAC (Abcam), PABP (Cell Signaling), GFP (Abcam) were purchased from the indicated vendors.

### Western blotting

Cells were lysed in RIPA buffer (50mM Tris pH8, 150mM NaCl, 1mM EDTA, 1% Triton X-100, 0.1% SDS) and protein concentrations determined by BCA assay (Pierce). Samples were then mixed with SDS sample loading buffer. We and others (Eoin Leen, personal communication) have observed that heating samples containing NS6, or some of the other NS6-containing precursors can cause these to precipitate in the sample buffer. As such blotting was performed by mixing the sample with SDS sample loading buffer and then immediately loading the sample onto the gel without heating. Samples were supplemented with RNase cocktail, and the use of lower sample concentrations (3-5μg/well) also helped prevent smearing. Samples were run on SDS-PAGE gels, transferred to nitrocellulose membranes, blocked and probed by standard protocols. Detection was accomplished using IR-dye 680 or 800-conjugated secondary antibodies followed by imaging on a Li-Cor Odyssey imager.

### Confocal Microscopy

Cells were grown on glass coverslips, transfected, and infected as indicated above. At the relevant timepoint, the coverslips were washed with cold PBS, fixed in 4% paraformaldehyde in PBS for 10 min on ice, and permeabilized with PBS containing 0.2% Triton X-100 and 4% paraformaldehyde for an additional 10 min at room temperature. Aldehyde groups were quenched with 0.2 M glycine in PBS. Subsequent antibody incubations and washing steps followed standard protocols and have been described previously (47). Alexa-fluor fluorescent secondary antibodies were obtained from Molecular Probes and used at the manufacturers recommended dilution. Coverslips were mounted on glass slides using Mowiol supplemented with DAPI. Imaging was performed on a Leica Sp5 confocal microscope using a 63x oil objective. Image analysis was performed in the Leica Lite software (Leica Microsystems).

### *In vitro* translation reactions

*In vitro* translation assays were performed with 1ug template plasmid and the T7 coupled *in vitro* transcription-translation kit (Promega). Reactions were either performed with ^35^S-methionine labeling, or unlabeled as indicated in the text. Half-size reactions (25μl) reactions were incubated at 30°C for 1h 30m. At the end of the incubation the reactions were mixed with an equal volume of Trans-stop buffer (10mM EDTA, 100ng/ml RNAse A) and incubated at room temperature for 30m.

For experiments assessing cleavage of ^35^S-methionine labeled substrate by unlabeled protease/precursors, after addition of trans-stop, 5μl of labeled substrate was mixed with 15μl of unlabeled protease. This was then mixed with 4 volumes (80μl) of protease assay buffer (10mM HEPES pH7.6, 0.1% CHAPS, 10mM DTT and 30% glycerol (46)) and incubated at 37°C for 48 hours.

All samples were then combined with reducing SDS-PAGE loading buffer, resolved on a denaturing 17.5% SDS-PAGE gel, dried, and imaged using a Typhoon FLA 7000 phosphorimager.

### LUMIER assay

All assays were performed in quadruplicate. 96 well plates containing 5 x 10^4^ HEK-293T cells per well in 100μl of DMEM supplemented with 15% FBS were seeded 12h prior transfection. Co-transfections were performed with 60ng of combined bait and prey plasmids with 0.3μl lipofectamine 2000 (Invitrogen) in 30μl Opti-MEM I (Gibco) per well. After 24h incubation, cells were lysed for 10 min at 4°C with shaking using 30μl of ice-cold LUMIER lysis buffer (PBS pH 7.4 (Mg/Ca free), 1% Triton X100, 0.1% BSA, 1mM DTT) with 1% Halt EDTA-free Protease/Phosphatase Inhibitor Cocktail (Pierce) and benzonase nuclease (Novagen)) Each supernatant was transferred in a 96 well V-bottom plate, and then centrifuged at 2000 x *g* for 15 min at 4°C. 20μl of each supernatant was retained and transferred to a new white flat bottom plate.

Dynabeads M280 sheep anti-rabbit IgG-coated magnetic beads (Invitrogen) were washed 3 times with the LUMIER washing buffer (PBS pH 7.4 (Mg/Ca free), 0.1% BSA, 1mM DTT, 1mM ATP) then coupled overnight at 4°C with a polyclonal anti-protein A rabbit antibody (Sigma-Aldrich). The beads were washed 4 times with LUMIER washing buffer. 2μl of beads in 30μl of LUMIER washing buffer was added to each well containing the lysate supernatents.

After 90 min of incubation on shaker at 4°C, 60μl of LUMIER washing buffer was added to each plate and mixed. After transferring 10μl of supernatant per well to new 96-well white plates plates to measure unbound activity, IP plates were washed six times with LUMIER washing buffer. Renilla luciferase activity was measured in both IP and unbound plates on a GloMax luminometer (Promega). Data were normalised to background ratios and Robust Z-score (RZ) calculated (48).

## Acknowledgements

IG is supported by a Wellcome Trust Senior Fellowship (WT097997MA). This work was also supported by the Cambridge NIHR BRC cell phenotyping hub, who are thanked for their assistance with microscopy.

---

**Figure S1.**
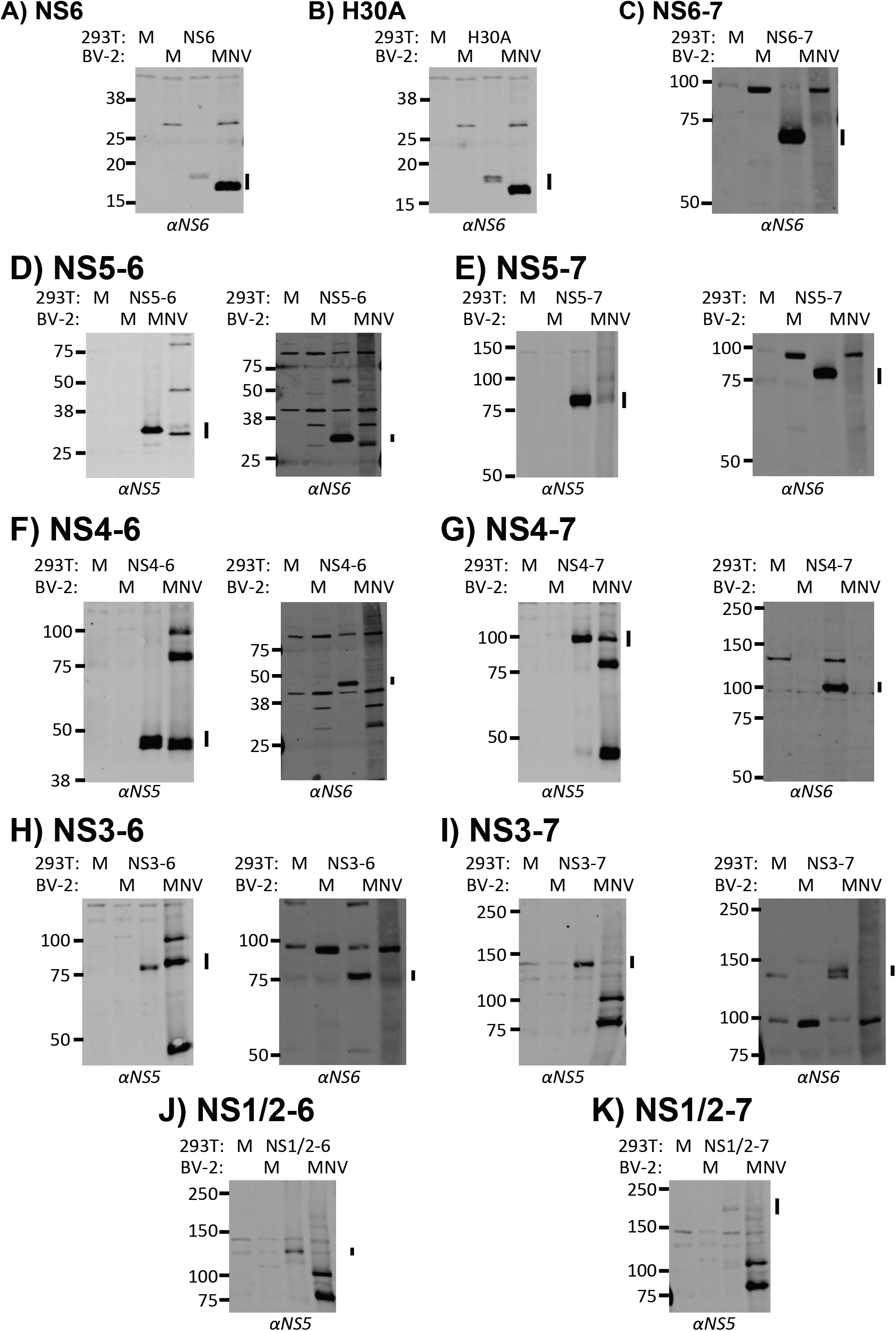
Confirmation of precursor identity in MNV-infected cell lysates. Lysates from BV-2 cells mock- or infected with MNV at MOI 10 and harvested at 9h post-infection, were compared to HEK-293T cell lysates mock- or transfected with plasmids expressing stable forms of the various precursors. Precursors were visualized by anti-NS6 and anti-NS5 antisera to assess co-migration. Note that the addition of a FLAG-tag on the precursors adds at additional 1.14kDa which results in some discrepancy in the migration of the smaller precursors.

**Figure S2.**
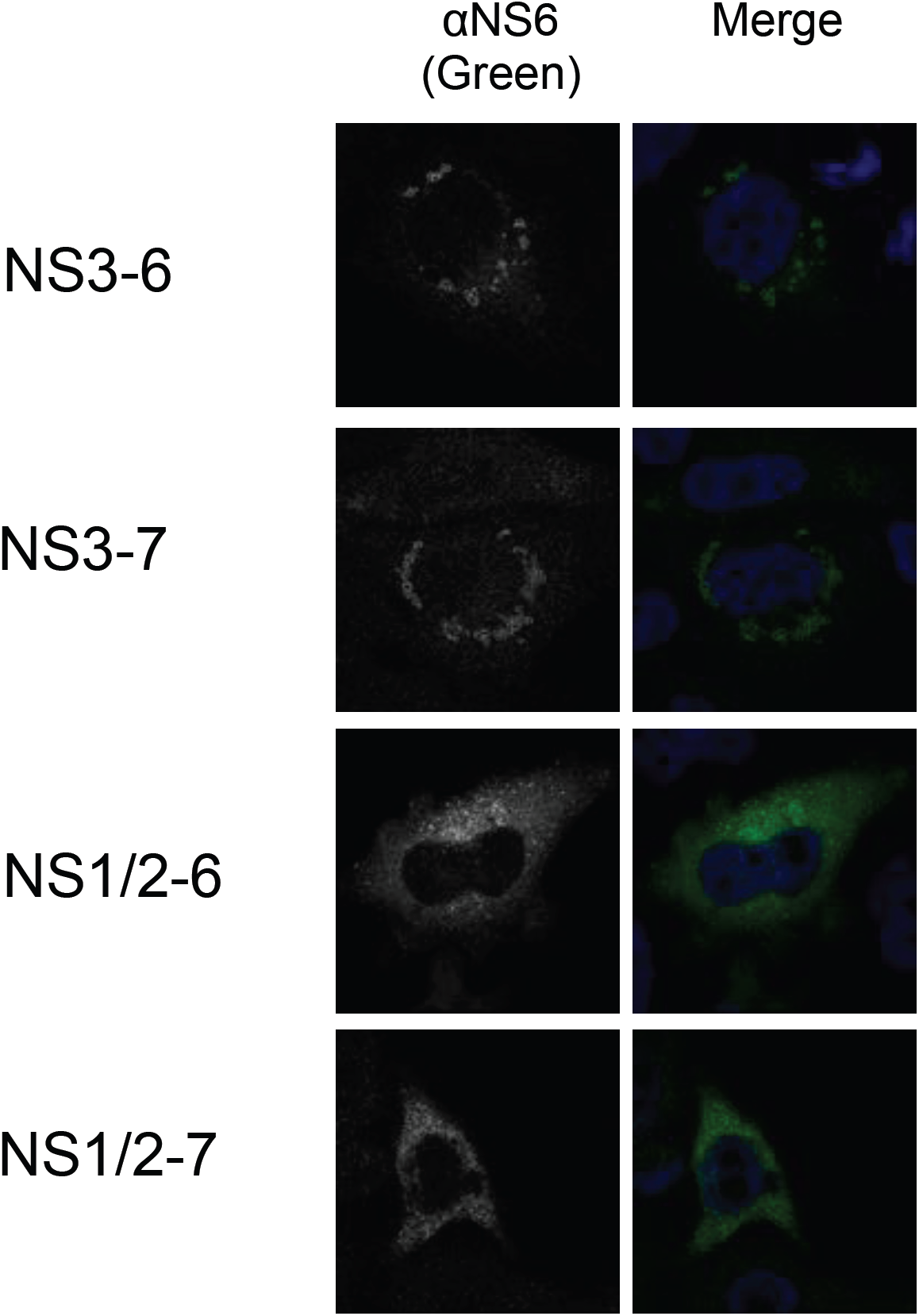
Confocal microscopy of N-terminal norovirus protease precursors. HeLa-CD300lf cells were transfected with plasmids expressing NS3-6, NS3-7, NS1/2-6 and NS1/2-7. At 18h post-transfection the cells were harvested and precursors visualized by anti-NS6 staining. Nuclei were visualized with DAPI.

**Figure S3.**
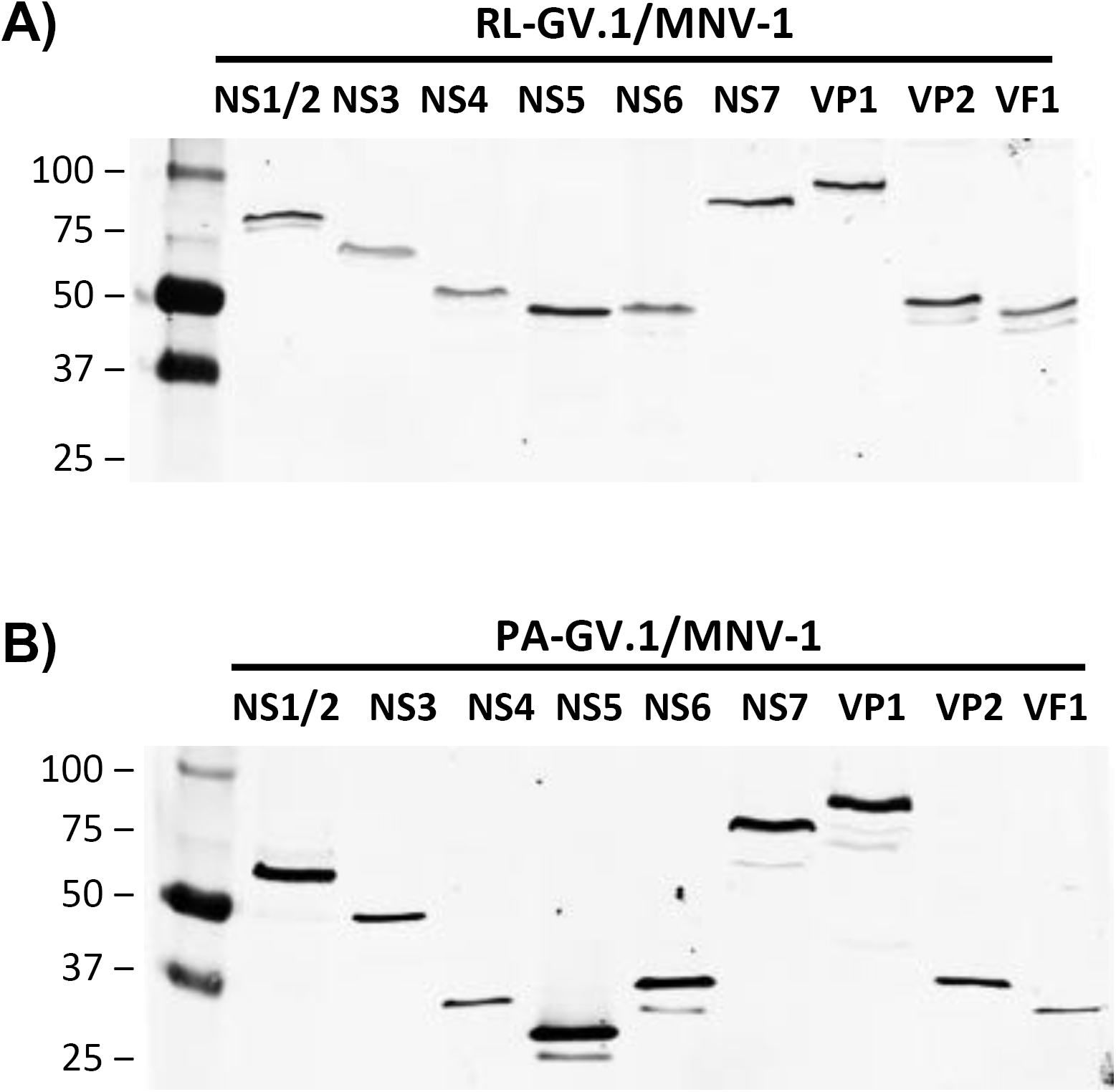
Western blotting of MNV LUMIER assay reagents confirms expression. Western blotting of lysates prepared from HEK-293T cells transfected with A) Renilla-luciferase or B) Protein-A fusions of the various MNV proteins confirms their expression and migration at the expected size. Samples were harvested at 24h post-transfection.

**Figure S4.**
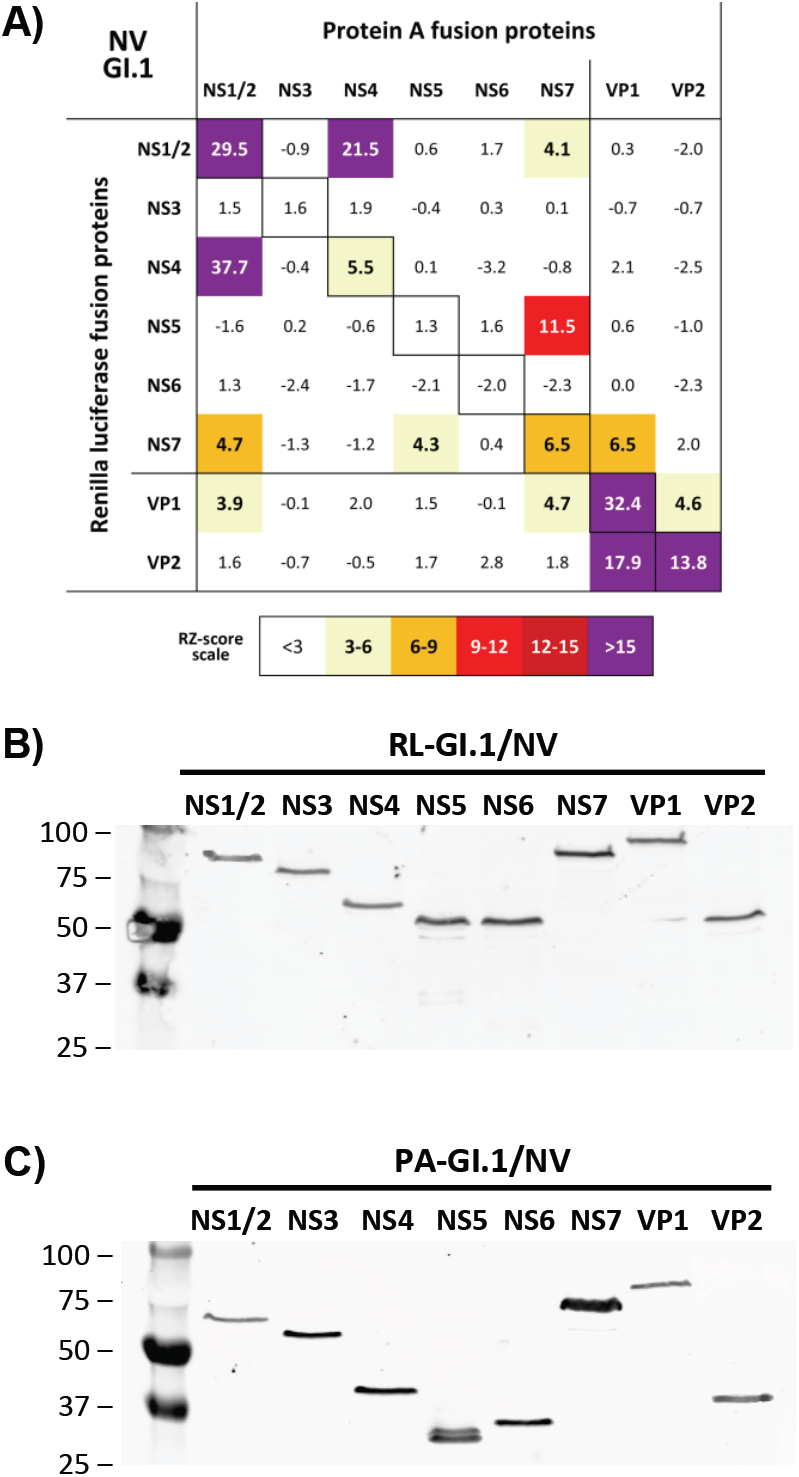
LUMIER assay reveals GI.1 replication complex interactions are conserved between GI.1 and MNV. A) HEK-293T cells expressing Protein-A and Renilla luciferase fusions of the various GI.1 human norovirus proteins were used for LUMIER analysis to identify protein:protein interactions. The numbers are robust z-scores. Positive protein:protein interactions are coloured by the strength of interaction with weak interactions showing in pale yellow, with the strongest interactions in purple. Western blotting of lysates prepared from HEK-293T cells transfected with B) Renilla-luciferase or C) Protein-A fusions of the various GI.1 norovirus proteins confirms their expression and migration at the expected size. Samples were harvested at 24h post-transfection.

**Figure S5.**
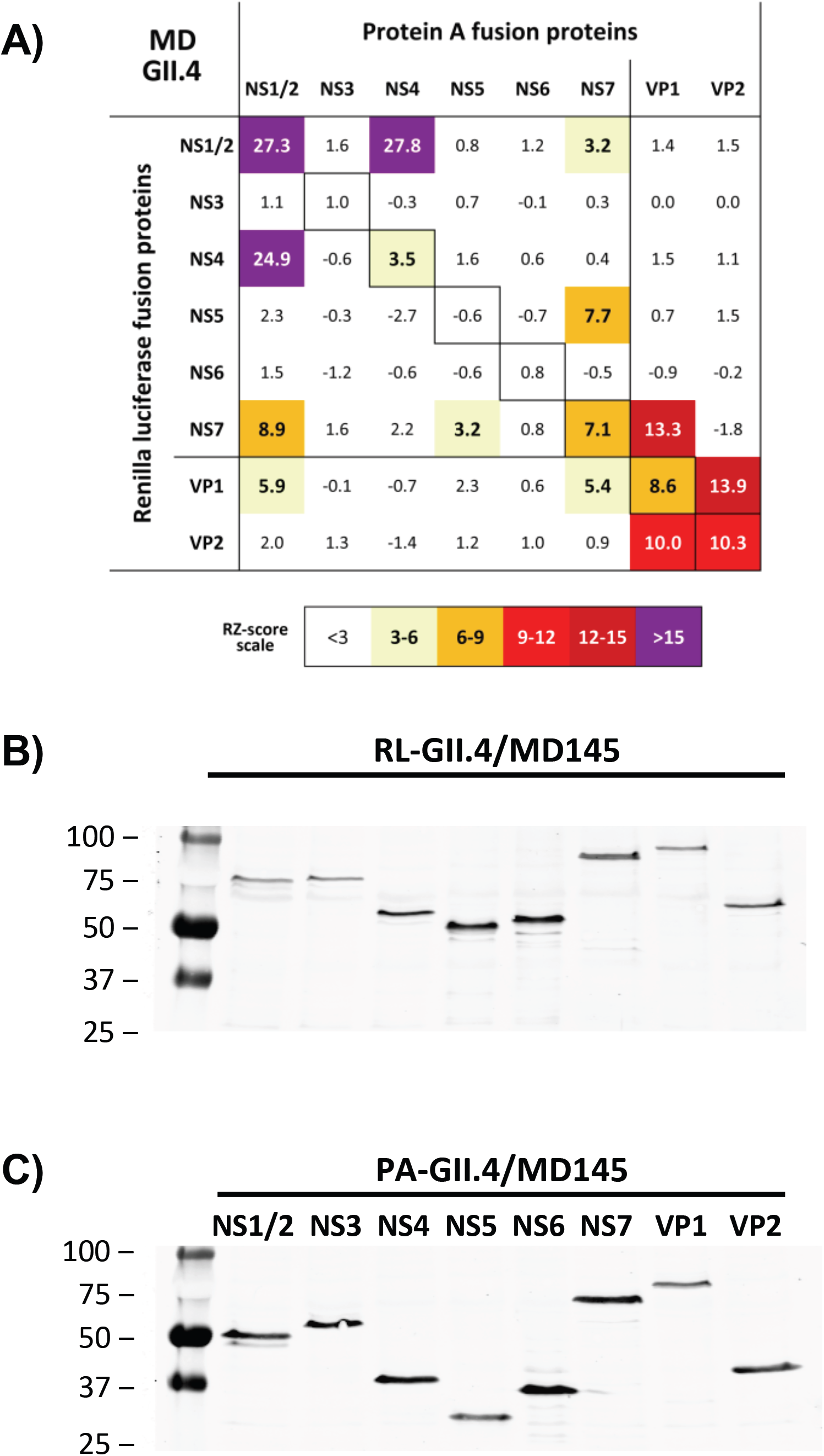
LUMIER assay reveals GII.4 replication complex interactions are conserved between GII.4 and MNV. A) HEK-293T cells expressing Protein-A and Renilla luciferase fusions of the various GII.4 human norovirus proteins were used for LUMIER analysis to identify protein:protein interactions. The numbers are robust z-scores. Positive protein:protein interactions are coloured by the strength of interaction with weak interactions showing in pale yellow, with the strongest interactions in purple. Western blotting of lysates prepared from HEK-293T cells transfected with B) Renilla-luciferase or C) Protein-A fusions of the various GII.4 norovirus proteins confirms their expression and migration at the expected size. Samples were harvested at 24h post-transfection.

**Table S1. Predicted molecular weights of norovirus proteins and precursor forms.** The first tab lists all the potential norovirus proteins and precursors that can be derived from the polyprotein. The second and third tabs list all the NS5- or NS6-containing precursors ranked by molecular weight from largest to smallest. Predicted molecular weight was calculated using the ExPASy server ProtParam tool.

## Supplemental Information

This manuscript has five supplementary figures and one supplementary table.

